# *Plasmodium caprae* infection is negatively correlated with infection of *Theileria ovis* and *Anaplasma ovis* in goats

**DOI:** 10.1101/515684

**Authors:** Hassan Hakimi, Ali Sarani, Mika Takeda, Osamu Kaneko, Masahito Asada

## Abstract

*Theileria, Babesia*, and *Anaplasma* are tick-borne pathogens affecting livestock industries worldwide. In this study, we surveyed the presence of *Babesia ovis, Theileria ovis, Theileria lestoquardi, Anaplasma ovis, Anaplasma phagocytophilum*, and *Anaplasma marginale* in 200 goats from 3 different districts in Sistan and Baluchestan province, Iran. Species-specific diagnostic PCR and sequence analysis revealed that 1.5%, 12.5%, and 80% of samples were positive for *T. lestoquardi, T. ovis*, and *A. ovis*, respectively. Co-infections of goats with up to 3 pathogens were seen in 22% of the samples. We observed a positive correlation between *A. ovis* and *T. ovis* infection. In addition, by analyzing the data with respect to *Plasmodium caprae* infection in these goats, a negative correlation was found between *P. caprae* and *A. ovis* and between *P. caprae* and *T. ovis*. This study contributes to understanding the epidemiology of vector-borne pathogens and their interplay in goats.

**Importance:** Tick-borne pathogens include economically important pathogens restricting livestock farming worldwide. In endemic areas livestock are exposed to different tick species carrying various pathogens which could result in co-infection with several tick-borne pathogens in a single host. The co-infection and interaction among pathogens are important in determining the outcome of disease. Little is known about pathogen interactions in the vector and the host. In this study, we show for the first time that co-infection of *P. caprae*, a mosquito transmitted pathogen, with *T. ovis* and *A. ovis*. Analysis of goat blood samples revealed a positive correlation between *A. ovis* and *T. ovis*. Moreover, a negative correlation was seen between *P. caprae*, a mosquito transmitted pathogen, and the tick-borne pathogens *T. ovis* or *A. ovis*.

## Introduction

Tick-borne diseases remain an economic burden for the livestock industry of tropical and subtropical regions of the world. Protozoan parasites such as *Babesia* spp. and *Theileria* spp. together with *Anaplasma* spp. are responsible for tick-borne diseases in small ruminants and cause great economic losses in the livestock-related industries (1, 2).

Small ruminant theileriosis is mainly caused by *Theileria lestoquardi, Theileria ovis*, and *Theileria separata. T. lestoquardi* is the most virulent specie and occasionally causes death while *T. ovis* and *T. separata* are benign and cause subclinical infections in small ruminants (3). Several species of *Babesia* have been described to cause ovine and caprine babesiosis including *Babesia ovis, Babesia motasi, Babesia crassa*, and *Babesia foliata* (4). *B. ovis* is the most pathogenic and causes fever, icterus, hemoglobinuria, severe anemia, and occasional death (5). *Anaplasma* spp. are important for human and animal health and these pathogens are generally considered to produce mild clinical symptoms. Although several *Anaplasma* spp. including *Anaplasma marginale, Anaplasma ovis*, and *Anaplasma phagocytophilum* could be found in small ruminants, *A. ovis* is the main cause of small ruminant anaplasmosis in the world. In Iran infections are usually asymptomatic and can produce acute symptoms if associated with stress factors (6, 7).

In Iran small ruminant farming is widely practiced with 52 and 26 million heads of sheep and goats, respectively, being raised mainly by small farmers (8). In regions with harsh and severe environments, such as central and southeast Iran, goat raising dominates. The great diversity of the environment in Iran affects the distribution of ticks, and thereby the pathogens transmitted. Several epidemiological studies are available regarding tick-borne pathogens in small ruminants in Northern and Western regions in Iran (9-12). However, there is a scarcity of data regarding the prevalence of *Babesia, Theileria*, and *Anaplasma* spp. infecting small ruminants in southeastern Iran. This study investigated the prevalence of tick-borne pathogens in Sistan and Baluchestan province, in the southeastern part of Iran where it borders with Afghanistan and Pakistan; and where the frequent border-crossing animal passage facilitates the circulation of tick-borne pathogens between countries (13).

In endemic areas livestock are bitten by vectors carrying multiple pathogens or different vectors transmitting various pathogens, which result in the development of multiple infections in the host. The interaction among different pathogens within a host are complex and may result in protection against virulent pathogens or exacerbate the clinical symptoms (14). In a study done on the indigenous African cattle in Kenya, it was shown that co-infection with less pathogenic *Theileria* spp. in calves results in a mortality decrease associated with virulent *T. parva* which is likely the result of cross protection (15). A recent study also showed a negative interaction between *B. ovis* and *T. ovis* in sheep, indicating infection with the less pathogenic *T. ovis* produces protection against highly pathogenic *B. ovis* (16). In contrast, co-infection of *B. ovata* and *T. orientalis*, two parasites which are transmitted via the same tick species, exacerbate the symptoms and produce clinical anemia in cattle (17). Recently we identified the goat malaria parasite, *Plasmodium caprae*, in goat samples originating from Sistan and Baluchestan province, Iran (18). However, nothing is known for this pathogen except some DNA sequence and morphology, and thus in this study we examined the prevalence of tick-borne piroplasms and *Anaplasma* spp. in these goat samples and evaluated the interplay among the identified pathogens.

## Results and Discussion

### Prevalence of *T. lestoquardi, A. ovis, B. ovis* and *Anaplasma* spp. in goat blood samples from Sistan and Baluchestan province, Iran

All 200 samples analyzed were negative for *B. ovis, A. marginale*, and *A. phagocytophilum;* while 3 (1.5%), 25 (12.5%), and 160 (80%) were positive for *T. lestoquardi, T. ovis*, and *A. ovis*, respectively (Table 2). In Zabol, 19/51 (37.3%) and 50/51 (98%) samples were positive for *T. ovis* and *A. ovis*, respectively, while *T. lestoquardi* was not detected. In Sarbaz, 3/125 (2.4%), 6/125 (4.8%) and 87/125 (69.6%) samples were positive for *T. lestoquardi, T. ovis* and *A. ovis*, respectively. None of the samples from Chabahar were positive for *T. lestoquardi* and *T. ovis* while 23/24 (95.8%) were positive for *A. ovis*. Sequence analysis of obtained PCR products confirmed that the species identities judged by PCR diagnosis were correct.

**Table 1.**
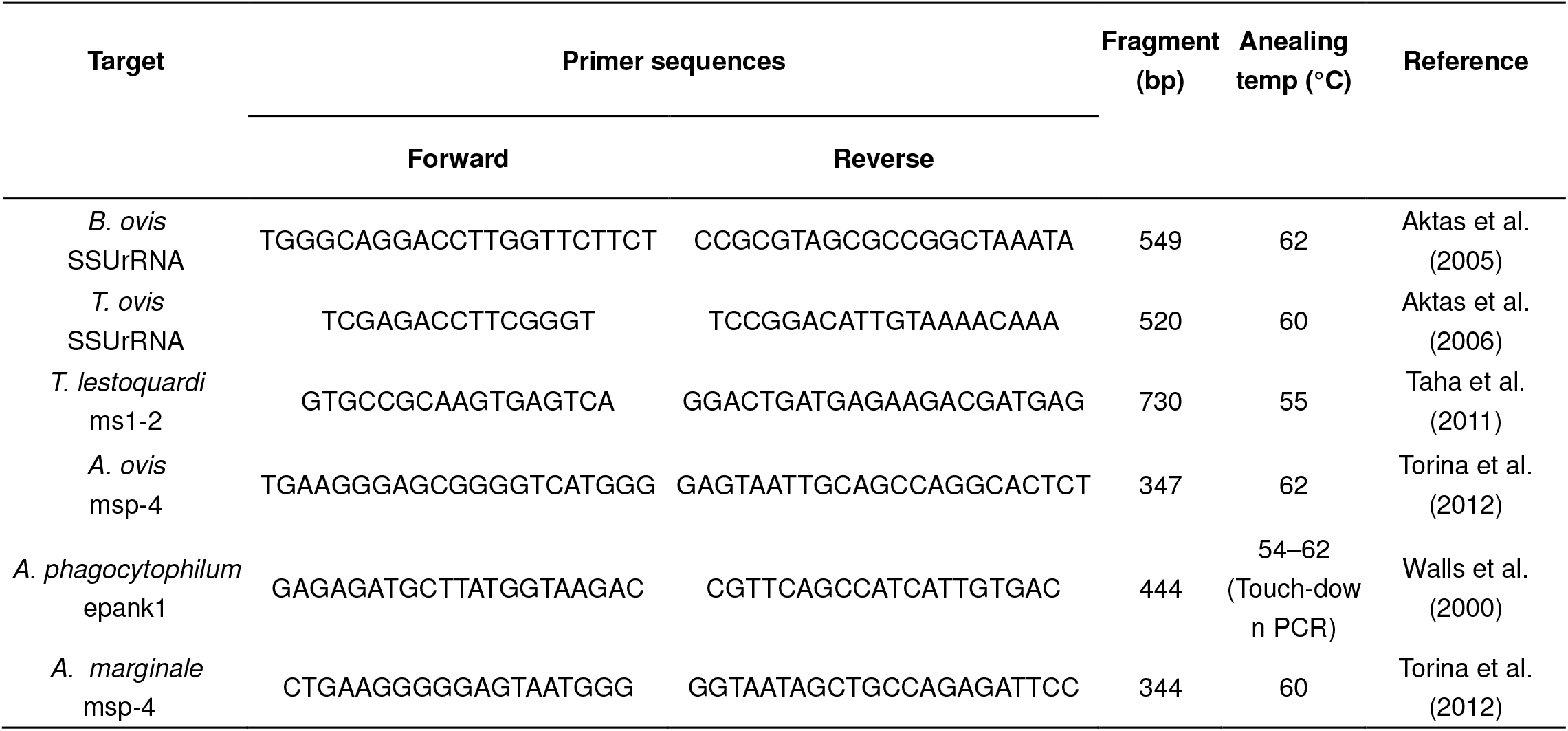
List of primers used in this study

**Table 2.** Pathogens identified in different districts in Sistan and Baluchestan

Hakimi et al., Table 2 Pathogens identified in different districts in Sistan and Baluchestan
| District | Pathogens |  |  | Total samples |
| --- | --- | --- | --- | --- |
|  | *T. ovis* | *T. lestoquardi* | *A. ovis* |  |
| Zabol | 19 (37.3%) | 0 | 50 (98%) | 51 |
| Sarbaz | 6 (4.8%) | 3 (2.4%) | 87 (69.6%) | 125 |
| Chabahar | 0 | 0 | 23 (95.8%) | 24 |
| Total | 25 (12.5%) | 3 (1.5%) | 160 (80%) | 200 |

Several species of *Theileria* can infect small ruminants and *T. ovis* and *T. lestoquardi* were reported previously from Iran (10, 19, 20). While there is no report on *T. ovis* prevalence in the goat population in Iran, the prevalence of *T. lestoquardi* is 6.25% and 19% in West Azerbaijan and Kurdistan provinces, respectively, in western Iran (21, 22). The prevalence of *T. lestoquardi* in sheep ranges from 6.6% in Razavi Khorasan province in northeast Iran to 33% in Fars province in central Iran, which is one of the most important endemic regions for ovine theileriosis in Iran (10, 23). *T. ovis* is more prevalent in sheep and ranges from 13.2% in western Iran to 73% in central Iran (10, 23). The overall infection rate of *T. lestoquardi* in this study was 1.5%, which is relatively low compared to the previous reports from Iran (21, 22). This difference may originate from the difference in sampling time, as all the positive samples in this study were collected in summer. The climate diversity which affects the distribution and infestation of the tick vector in various regions in Iran (24) may also contribute to a difference in infection rates. *T. ovis* was present in 12.5% of samples and this is the first molecular report of this parasite in goats in Iran. *B. ovis* is one of the most important and highly pathogenic parasites that infects small ruminants and prevalent in different regions in Iran (11). Although *Rhipicephalus bursa*, the tick vector of *B. ovis*, exists in Sistan and Baluchestan (24), we could not detect this pathogen in this study suggesting that *B. ovis* may not be common in the surveyed region.

Goat anaplasmosis in Iran is mainly caused by *A. ovis* and *A. marginale* (12). The infection rate of *A. ovis* is from 34.7% in western Iran to 63.7% in northern and northeastern Iran (12, 25). The overall infection rate of *A. ovis* in goats was 80% in this study which was higher than other reported areas in Iran. The difference in sampling time, diagnosis method, geographical, and climate variation may contribute to the differential prevalence of this pathogen.

There is no epidemiological report on tick-borne pathogens in small ruminants in Afghanistan, and similar reports are limited from Pakistan and not from the western region (26, 27). Thus, the result of this study serves as a useful reference to estimate the prevalence of tick-born piroplasms and *Anaplasma* spp. in the Afghanistan and the Pakistan regions neighboring the Sistan and Baluchestan province of Iran.

### Interplay between *P. caprae* and other pathogens co-infected in the goat

Because the goat malaria parasite, *P. caprae*, was detected in 28 samples among these 200 samples in the previous study (18), we evaluated the possible effect of a specific pathogen infection against the other pathogens identified in this study. A correlation coefficient value (R_ij_) was calculated for each two-pathogen interaction (28). Co-infections are summarized in Table 3. A strong negative correlation between *P. caprae* and *A. ovis* infections with R_ij_ value of −0.593 was observed (p < 0.01 by Chi-square test, Table 4) and their double infection was only 8%. Infection of *P. caprae* and *T. ovis* showed a relatively weak negative correlation, yet significant (R_ij_ value: −0.182, p < 0.05 by two-tailed Fisher’s exact test). However, all *T. ovis* positive samples were also positive for *A. ovis* and a relatively weak, though significant, positive correlation was observed (R_ij_ value: 0.118, p < 0.01 by two-tailed Fisher’s exact test).

**Table 3.** Pathogens detected in goat samples from Sistan and Baluchestan

Pathogens detected in goat samples from Sistan and Baluchestan
| Pathogens | Positive numbers (%) |
| --- | --- |
| Single infection |  |
| *T. lestoquardi* | 1 (0.5) |
| *A. ovis* | 118 (59) |
| *P. caprae* * | 12 (6) |
| Double infection |  |
| *A. ovis* & *T. ovis* | 24 (12) |
| *A. ovis* & *T. lestoquardi* | 1 (0.5) |
| *A. ovis* & *P. caprae* | 16 (8) |
| Triple infection |  |
| *A. ovis*, *T. ovis* & *T. lestoquardi* | 1 (0.5) |

\*From Kaewthamasorn et al, 2018

**Table 4.** Correlation between pathogens

Correlation between pathogens
| Pathogens | *A. ovis* | *T. ovis* | *T. lestoquardi* |
| --- | --- | --- | --- |
| *P. caprae* | **-0.593 (0.0011)** | **-0.182 (0.029)** | -0.059 (1) |
| *A. ovis* | — | **0.118 (0.0055)** | -0.131 (0.49) |
| *T. ovis* | — | — | -0.072 (0.33) |

Correlation coefficient between two pathogens ( $R_{ij}$ ) is presented. The $p$ -value is shown in the bracket. $p$ -values were calculated by chi-square test (*P. caprae* and *A. ovis*) or Fisher's exact test (the other combinations).

The negative correlation between two vector-borne pathogens could also happen in the host or the vector by competing for resources such as space like available erythrocytes, nutrients, or impact on the immune response. We found a negative correlation between *P. caprae* and *A. ovis* and between *P. caprae* and *T. ovis*. Mosquitoes are the likely vector for *P. caprae*, while *A. ovis* and *T. ovis* are transmitted by ticks; thus excluding the possibility of negative interference in the vector and highlighting likely competition in the goat (29, 30). A negative association was reported between *Theileria annulata*, a protozoan parasite responsible for tropical theileriosis, and *A. marginale* (28). In a study that was done using blood samples from sick sheep, the authors showed that the presence of *T. ovis* was negatively correlated with *B. ovis*, indicating that infection with low pathogenic *T. ovis* protects sheep from infection with highly pathogenic *B. ovis* (16). An absolute exclusion was shown to exist between *T. annulata* and *B. bovis*, since the authors did not find any co-infection in cattle samples in Algeria (28). The negative correlation between two pathogens could happen through modification of host immune response such as development of cross-protection immunity. Alternatively, this may be due to a mechanical interference between pathogens since all these pathogens infect host erythrocytes. However, there is no data on the erythrocyte type preference and receptors for these pathogens. Studies on the molecular mechanism of erythrocyte invasion and modification mediated by these pathogens would provide important insights behind these observations; however, such information are scarce, if any. Given the fact that *P. caprae* observed in the goats had very low parasitemia, below the microscopy detection limit (18), we consider that the interference by *P. caprae* against *A. ovis* and *T. ovis* is quite unlikely and exclusion may take place through modulating the host immune system or mechanical interference by *A. ovis* and *T. ovis*.

The best example of positive correlation among two vector-borne pathogens is between *Borrelia burgdorferi*, the causative agent of Lyme disease, and *Babesia microti*, the primary agent of human babesiosis, both of which are transmitted by the tick *Ixodes ricinus* (31). Co-infection of these two pathogens are common and enhance the transmission and emergence of *B. microti* in human population in USA, possibly by lowering the ecological threshold for establishment of *B. microti* (14, 32). Immunosuppression by one pathogen may predispose the host to the second pathogen. This phenomenon could be seen in *B. microti* with the parasites *Trypanosoma musculi* and *Trichuris muris* in mice (33, 34). Moreover, the possibility of coinfection increases if the pathogens are transmitted by the same vector (34). Both *T. ovis* and *A. ovis* are transmitted by the same tick, *R. sanguineus*, in the region thus positive correlation may be a result of the simultaneous inoculation of these pathogens to the goat by ticks or by enhancing pathogen fitness and transmission by tick vector (30, 35). In addition, positive correlation of *T. ovis* and *A. ovis* infection may suggests the absence of cross protection between these pathogens; one eukaryotic protozoan parasite and the other prokaryotic bacteria. One infection appears to increase the susceptibility to the other pathogen. Co-infection is often associated with exacerbation of symptoms, thus, competition among pathogens could be beneficial to the host (17). It is worth investigating the mechanism for competitive interaction among these pathogens.

The distribution of *T. lestoquardi, T. ovis*, and *A. ovis* in different regions in Iran is well reported. However, in this study we focused in Southeast of Iran, Sistan and Baluchestan, where no reports exist, and showed the co-infection of these pathogens. Coinfection of several pathogens might influence the pathogenesis in the host and may jeopardize correct diagnoses. We showed a negative correlation between *A. ovis* and *P. caprae* and between *T. ovis* and *P. caprae*, suggesting possible interference via immunity or against erythrocyte invasion by the other pathogen. The results of this study may contribute to understand these pathogen interactions in the host, and aid in designing preventive measures of tick-borne pathogens in the region.

## Materials and Methods

### Sampling sites and blood collection

Blood samples were collected from 200 goats from 3 districts in Sistan and Baluchestan provinces, including Zabol (n = 51), Sarbaz (n = 125), and Chabahar (n = 24) as shown in Fig. 1. In each district, blood samples were collected from different farms. Sampling was done in January, June, and November of 2016 and July of 2017. Blood sampling and DNA extraction was performed as described (18).

**Fig 1.**
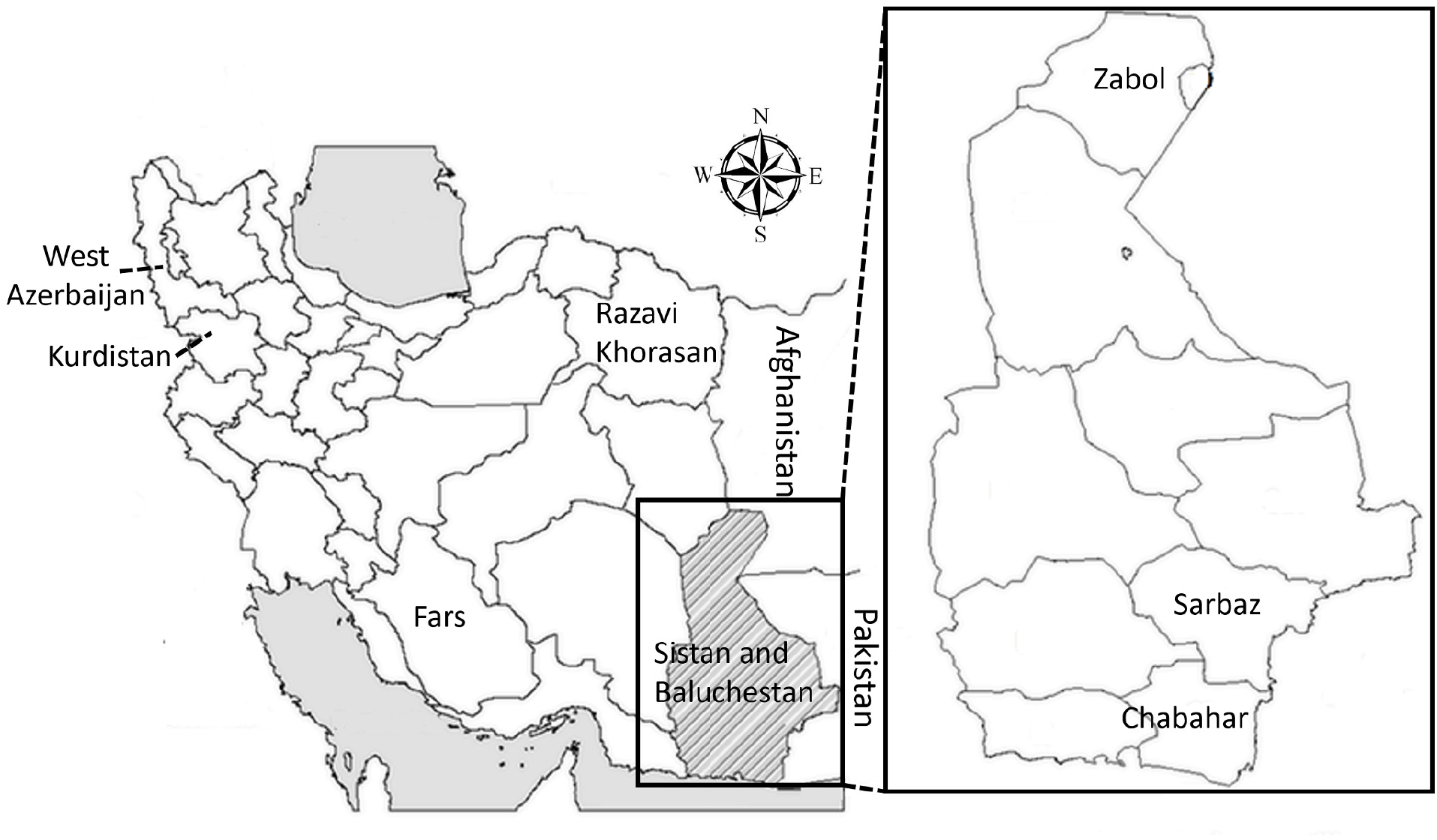
Map of Iran showing sampling sites in Sistan and Baluchestan province, Iran.

### Ethical statement

Sampling of goats was performed with the informed consent of the farm owners. All procedures were carried out in compliance to ethical guidelines for the usage of animal samples permitted by University of Zabol (IRUOZ.ECRA.2016.01).

### Detection of *Babesia* spp. *Theileria* spp. and *Anaplasma* spp. by species-specific PCR

Each DNA sample was screened for *B. ovis, T. lestoquardi, T. ovis, A. phagocytophilum, A. ovis*, and *A. marginale* by species-specific PCR as described (36–40). Small subunit ribosomal RNA (SSUrRNA) was the target gene for detection of *B. ovis* and *T. ovis* while specific primers targeting merozoite surface antigen gene *(ms1)* were used for detection of *T. lestoquardi. A. ovis*, and *A. marginale* were screened using primers targeting major surface protein 4 gene (*msp-4*); and for specific detection of *A. phagocytophilum*, primers targeting *epank1* gene were used (Table 1). A correlation coefficient (R_ij_) between the different pairs of pathogens was measured as described (28). The 27 negative goats were excluded from the calculation of correlation coefficient. Chi-square text and two-tailed Fisher’s exact test were used to evaluate the significance of association between co-infected pathogens.

### Cloning and sequencing

PCR products of all positive samples for *T. ovis* or *T. lestoquardi* and three positive samples for *A. ovis* randomly selected from each region were sequenced. The amplified PCR products were recovered from agarose gels and cloned into the Zero Blunt TOPO vector (Thermo Fisher Scientific, Carlsbad, CA, USA) according to the manufacturer’s protocol. Following transformation three *E. coli* colonies were selected, the plasmids were extracted and purified, and the gene sequences were analyzed using BigDye Terminator v1.1 and an ABI 3730 DNA Analyzer (Applied Biosystems, CA, USA). The single nucleotide polymorphisms (SNPs) found in the obtained sequence were confirmed by repeating the amplification, cloning, and sequencing process. *T. lestoquardi ms1* (*Tlms1*), *T. ovis* SSUrRNA (ToSSUrRNA), and *A. ovis msp-4* (*Aomsp-4*) sequences from this study were deposited in GenBank (*T. ovis*: LC430938 and LC430939, *A. ovis:* LC430940, LC430941 and LC430942, *T. lestoquardi:* LC430943, LC430944, LC430945, LC430946, LC430947 and LC430948).

## Acknowledgments

H.H. is a recipient of the JSPS Postdoctoral Fellowship for foreign researchers from the Japan Society for the Promotion of Science. This study was supported in part by JSPS grants-in-Aids for Scientific Research (B), No 16H05807 to MA and OK, and Scientific Research (C), No 16K08021 to MA. The funders had no role in study design, data collection and interpretation.

The authors are grateful to Paul Frank Adjou Moumouni and Xuenan Xuan from the National Research Center for Protozoan Diseases, Obihiro University of Agriculture and Veterinary Medicine for providing positive control DNA for performing PCR. We thank Thomas J. Templeton from our institute for his critical reading of the manuscript.

